# A simple and accurate method for inferring missing ploidy information from sequence data

**DOI:** 10.64898/2026.09.06.749761

**Authors:** Shruti V. Kulkarni, Andrew A. Crowl, George P. Tiley

## Abstract

Polyploidy can be a critical factor for explaining plant trait variation, niche diversification, or speciation. However, inferring ploidy from silica-dried or historical samples using chromosome counts or flow cytometry is not possible, and scaling up ploidy estimation to population-level fresh contemporary samples can be challenging as well. Thus, we present a new method for estimating ploidy levels directly from sequencing data using machine learning – the Polyploid Population Genomics Tool Kit (PPGTK). The machine-learning approach is advantageous as it relaxes the assumptions of previous probabilistic methods and provides per-sample probabilities, allowing investigators to evaluate uncertainty in their system of interest.. We demonstrate performance and accuracy of the method on simulated and empirical data. Simulations showed above 99% accuracy, even for low coverage data, as long reads were mappable to the reference genome. For empirical analyses, we used target enrichment data from blueberry wild relatives (*Vaccinium* sect. *Cyanococcus*) and whole-genome data from sweetpotato wild relatives (*Ipomoea* ser. *Batatas*). Ploidy was recovered with 99% accuracy across 70 *Vaccinium* individuals and 97% across 82 *Ipomoea* individuals. Analysis of many individuals is fast and requires only a multisample VCF, which is presumably generated for the research anyway, and some samples of known ploidy for training the classifier. The approach implemented in PPGTK is promising for collections-based research as well, enabling ploidy classification of historical specimens based on present-day observations. The method is implemented in a new Python package as a single command that can run on a conventional laptop.

## INTRODUCTION

Polyploidy is a pervasive and recurrent feature of plant genome evolution, and variation in ploidy within and among species shapes reproductive isolation, ecological breadth, and adaptive potential (e.g., Stebbins 1985; Heslop-Harrison et al. 2021). Interpreting that variation requires ploidy data for individuals rather than for species alone, as mixed-ploidy populations are common and cytotype can track niche and geography rather than taxonomy (Crowl et al. 2017; Baniaga et al. 2020; Gaynor et al. 2025; Phillips et al. 2025). The gold-standard approaches — chromosome counts and flow cytometry (Pellicer et al. 2021) — require fresh or specially preserved tissue and do not readily scale to the hundreds of individuals that population genomic studies now sequence routinely. This often creates a discrepancy, in which genomic data are generated for a large number of samples while ploidy is known for only a subset. Inferring ploidy directly from sequence data would close the gap at negligible cost and extend ploidy inference to material for which cytological methods are intractable, including museum specimens.

Several methods have been introduced to estimate ploidy directly from genetic or genomic data. At the most basic level, counting the number of unique haplotypes or the maximum number of alleles observed from fragments such as microsatellite loci has been used (Gompert and Mock 2017; Hagl et al. 2021). Such counting methods suffer from concern regarding allele dropout, sampling error, and the need to identify informative loci (e.g. Gompert and Mock 2017; Seeber et al. 2014). Genomic data have allowed a new wave of effective k-mer-based methods (Ranallo-Benavidez et al. 2020; Rhie et al. 2020). K-mer approaches leave a gap between fragments and whole genomes though, such as subsampled portions of the genome generated by restriction digests (e.g. RADseq) or probes that enrich targeted loci (e.g. Hyb-Seq). A number of approaches based on allele balance data, or the fraction of reads that support an alternate allele with respect to a reference, have been suggested based on mixture modeling (Weiß et al. 2018; Tiley et al. 2024; Gaynor et al. 2024). However, complications from model over-fitting are a concern and false positives would plague most studies of natural populations without extensive validation and manual curation. One preceding study also used allele balance (AB) data, but discretized the distribution into bins and used those bins as features for a principal component analysis and subsequent clustering (Gompert and Mock 2017). Assignment of individual sites to bins was determined with a probabilistic Bayesian model to explore all possible genotypes, which can be challenging in terms of compute burden and practical implementation for large numbers of individuals.

Here, we build upon the conceptual approach of Gompert and Mock (2017), but simplify the procedure using machine learning to facilitate classification of unknown samples given some known individuals. The method works with a multisample variant call format (VCF) file regardless of the underlying data type (e.g. target enrichment, RADseq, whole-genome). We explored multiple supervised learning strategies, but found that a straightforward logistic regression approach performed well in both simulations and empirical validation datasets. For empirical analyses, we re-analyze Hyb-Seq data from blueberries and their wild relatives, *Vaccinium* sect. *Cyanococcus* (Crowl et al. 2022; Manos et al. 2026) as well as whole-genome data from sweetpotatoes and their wild relatives, *Ipomoea* ser. *Batatas* (Wu et al. 2018; Muñoz-Rodríguez et al. 2022; Yan et al. 2024). Both cases represent complexes of diploids, tetraploids, and hexaploids where morphological determination of ploidy is difficult and the evolutionary history of economically important species remains uncertain. The ploidy classification method has been implemented through a Python package, the Polyploid Population Genomics Tool Kit (PPGTK) and is available on GitHub (https://github.com/tileylab/PPGTK). The function *classify-ploidy* only requires only a multisample VCF file and a metadata CSV file that minimally includes sample names and some known ploidy levels (Fig. 1).

**Figure 1.**
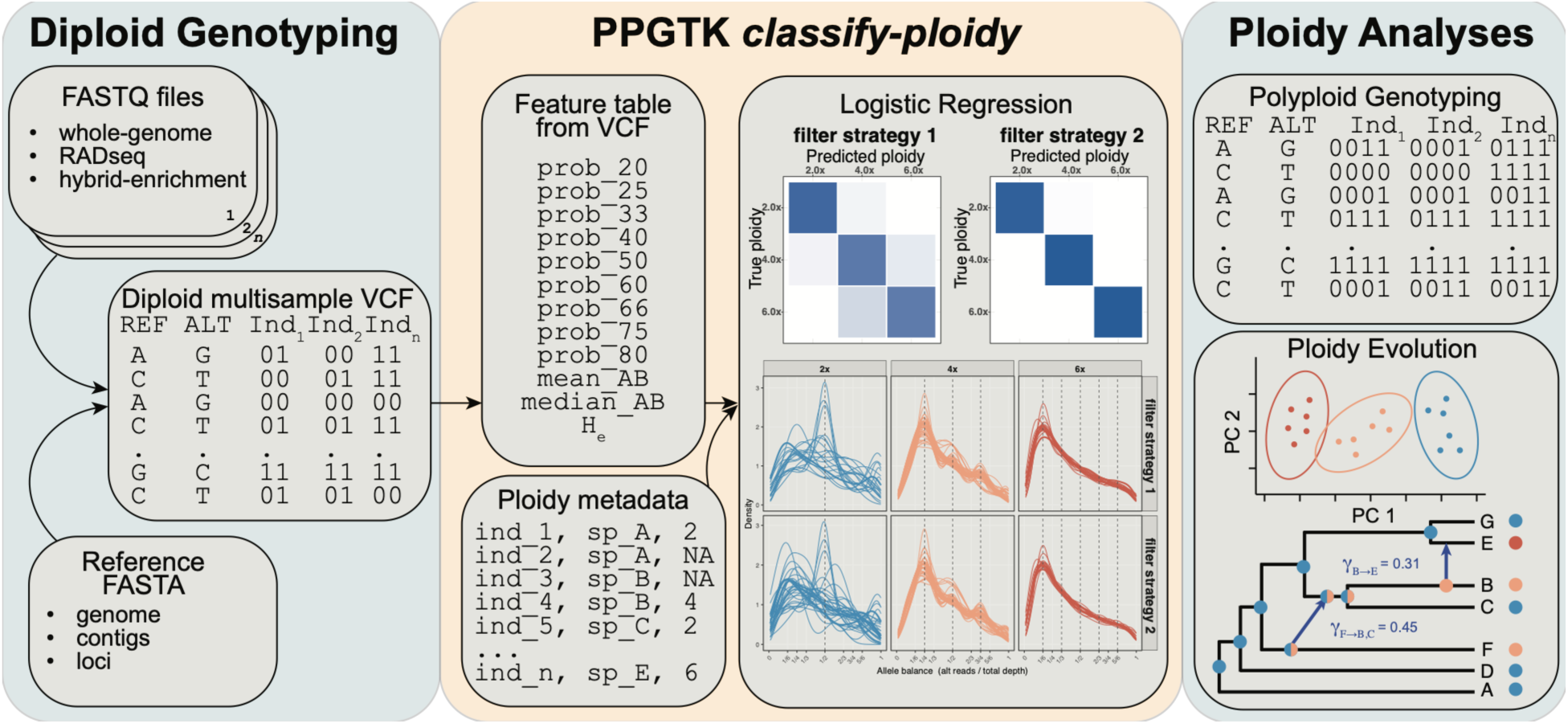
Overview of inputs and features used by *classify-ploidy*. Downstream analyses can leverage the predicted ploidies for genotyping or analysis.

## METHODS AND RESULTS

### Logistic regression classifier

To complement other ploidy estimation approaches, we implemented a simple machine-learning approach in Python using logistic regression and cross validation based on the scikit-learn library (Pedregosa et al. 2011). Although expectations can be developed for allele balance ratios at different ploidy levels, such expectations are difficult to meet in practice due to incorrect polarization of ancestral versus derived alleles (Keightly and Jackson 2018), reference bias in the genotyping process (Gerard 2018), and other artifacts that might arise from read stochasticity, error, and paralogy. A supervised approach was thus preferred, so a classifier can be trained on observations from known samples, and hopefully avoid false positives associated with overfitting from probabilistic mixture models characterized in previous methods (Weiß et al. 2018; Tiley et al. 2024; Gaynor et al. 2024) or ambiguity in higher ploidy levels (Viruel et al. 2019; Gaynor et al. 2024).

The method works from a multisample VCF, treating all individuals as diploid, that conforms with the v4.3 specification. The VCF v4.3 specification would be the product of any up-to-date joint-genotyping protocol, such as with GATK (McKenna et al. 2010). While a user can have as many annotations in the VCF as needed for filtering and analysis, PPGTK *classify-ploidy* only uses the allele depth (AD) field for biallelic variants. For each individual, variants passing filters are used to calculate per-site allele balance (AB), such that *AB* = 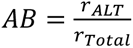 where *r_ALT_* is the number of reads supporting the alternate allele and *r_Total_* is the total number of reads supporting the reference or alternate allele. The AB distribution is then discretized into bins where each value is grouped to the closest bin at 0.2, 0.25, 0.33, 0.4, 0.5, 0.6, 0.66, 0.75, and 0.8. Although the bins are unbalanced, these AB values represent expected ratios under diploid, triploid, tetraploid, pentaploid, and hexaploid (partially) scenarios. Any higher ploidy levels are assumed to be sufficiently characterized by the bin distribution, and the naïve approach to discretization avoids computationally expensive enumeration of possible genotype likelihoods (Goempert and Mock 2017). Because direct counts would potentially bias inference by variation in sequencing depth, bin counts are standardized as proportions. The central tendencies of AB data are used for model training as well, such that the mean and median of the distribution are calculated per-individual. Global per-site heterozygosity is also calculated and used as a feature for model training, as increasing ploidy should generally increase observed heterozygosity. Thus, the genotype data is represented by a compact feature profile suitable for downstream machine-learning analyses.

For supervised ploidy classification, samples with available ploidy labels can be used to train a logistic regression model capable of predicting ploidy from the extracted AB and heterozygosity features. Although we initially explored multiple classifiers, logistic regression was chosen as the only method currently available in PPGTK as ploidy classification generally appeared a linearly separable problem while providing easy interpretation for end-users. Five-fold cross-validation is used by default to estimate accuracy and performance metrics on the known individuals to provide some measures of confidence in the predictions. For training data, standard performance metrics that evaluate correctness of the model are used: precision 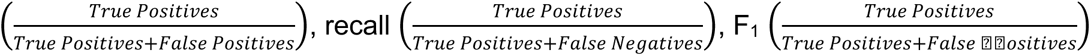, and 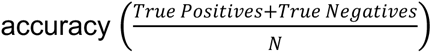 is reported based on all individuals as well as for each ploidy class to help identify weaknesses in the model. In the case of unequal training data, for example where diploids are much more prevalent than polyploids, the balanced accuracy metric can be used. For each individual, the predicted ploidy along with probabilities for each class are returned to the user. The probabilities can be used to distinguish high-confidence predictions from ambiguous cases that may require additional review, or additional data to resolve. To test the method performance, we use both simulations and two moderately-sized empirical datasets.

### Simulations

To explore accuracy of the method and its limitations, we performed simulations under the multispecies coalescent (MSC; Rannala and Yang 2003) model such that a tetraploid and hexaploid lineage were from contemporaneous progenitors, similar to Tiley et al. (2024). A fixed species topology (Fig. 2) was used and the nucleotide diversity parameter (*θ*) was constant over the tree. A *θ* of 0.0001, 0.001, or 0.01 was used and the lineage split times *τ* was scaled to *θ* such that the rate of evolution was the same across simulations, and only time differed. To provide context, assuming a per-generation mutation rate of *μ* = 10^-8^ and a generation time of one year, these *θ* values equate to a hybrid age of 50 thousand years ago (kya), 500 kya, and five million years ago (mya), respectively. Simulations were also performed for a *θ* of 0.0004, 0.004, and 0.04 to increase incomplete lineage sorting (ILS) or the overall rate of evolution while holding divergence times the same.

**Figure 2.**
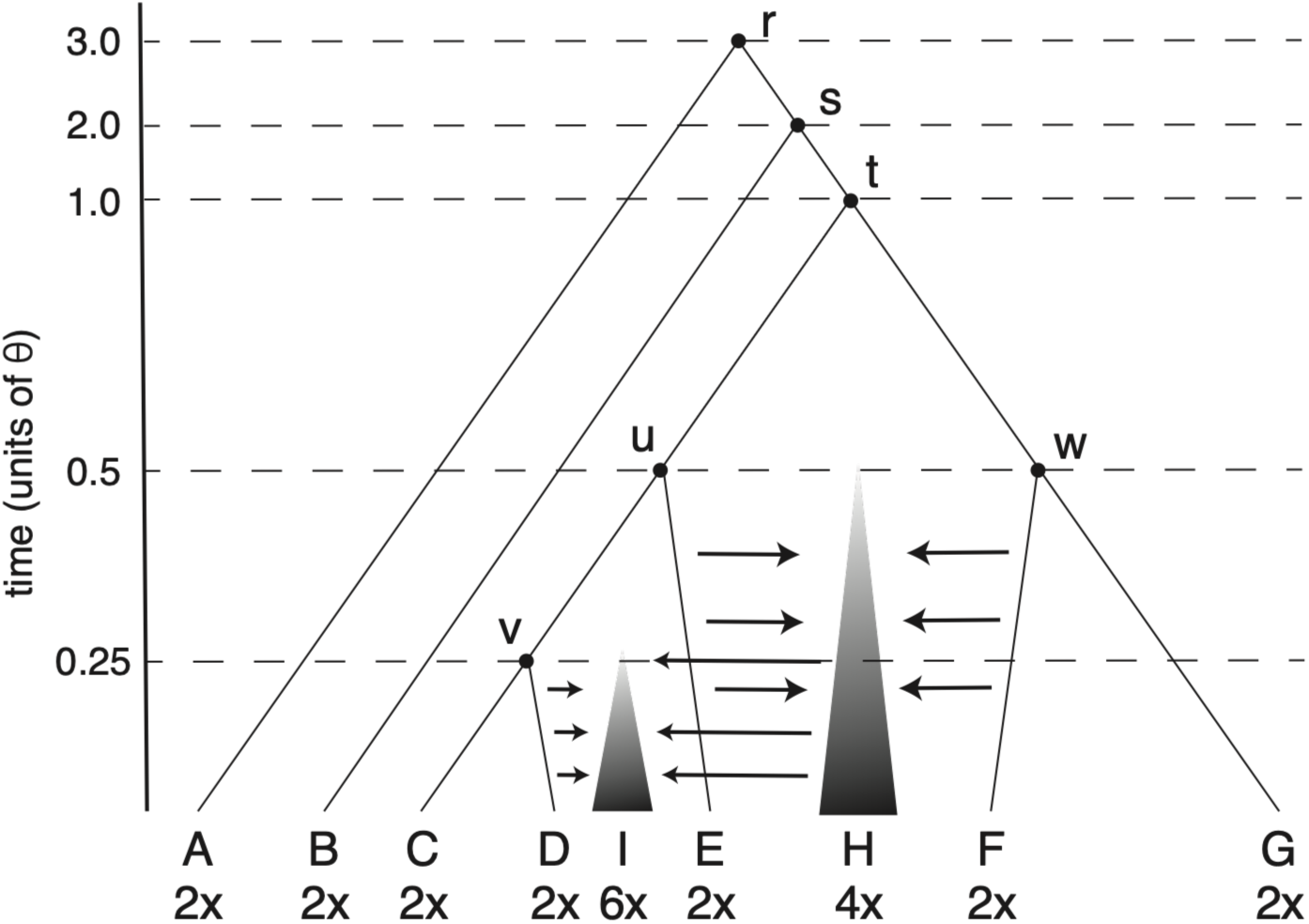
Tree used for simulation. The tetraploid species H and hexaploid species I are formed by randomly sampling haplotypes from their progenitors. Letters at nodes are simply node names used for setting up simulations. Time in units of population size reflect parameter scaling for simulations where *θ* is 0.0001, 0.001, or 0.01.

For each simulation condition, 1000 loci 500 base pairs (bps) in length were simulated 100 times. 40 haplotypes were simulated per locus per species.

For each diploid species, two haplotypes were chosen at random and used to simulate paired-end 150bp FASTQ data at 100x depth with wgsim at an error rate of 0.01 to mimic Q20 reads (Li and Durbin 2009). The reads were simulated proportionally from each haplotype, such that each haplotype has a depth of 50x. The 100x data was then randomly downsampled to depths of 10x, 20x, and 40x. The random downsampling does not guarantee proportional representation of haplotypes such that some read stochasticity would be possible at low depths. The polyploid species in the simulation re-used the same 100x read pools, but sampled randomly from each progenitor haplotype. Any tetraploid individual from species H can be traced to two haplotypes from E and two haplotypes from F, and any hexaploid individual from species I has six underlying haplotypes from species D, E, and F. This simulation and sampling strategy resulted in 20 individuals per species across all analyses, with 140 diploids, 20 tetraploids, and 20 hexaploids.

A simplified joint-genotyping pipeline was used to analyze simulated read data. For each simulation replicate, a reference genome was constructed by randomly sampling one haplotype per locus from species A. Simulated reads were then aligned to the reference genome with BWA v0.7.19 using the MEM algorithm (Li 2013). Genotyping then proceeded with GATK v4.6.2.0 (McKenna et al. 2010) HaplotypeCaller (Poplin et al. 2017) in GVCF mode, followed by CombineGVCFs and GenotypeGVCFs. The resulting multisample VCF from GenotypeGVCFs was then used for analysis without any additional filtering. The number of variants resulting from the simulation and genotyping process varied with simulation condition, but there was a notable drop in information for the simulation conditions with hybridization at five mya, where the reference genome was highly divergent (Table S1)

Ploidy classification was nearly perfect across simulation conditions where hybridization occurred at 50 or 500 kya for both standard and high ILS scenarios (Fig. 3; Table S2). Classification was not feasible for the high divergence, 5 mya hybridization, scenarios, because reads were no longer mappable to the reference genome (Supplementary Figs. S1-S6).

**Figure 3.**
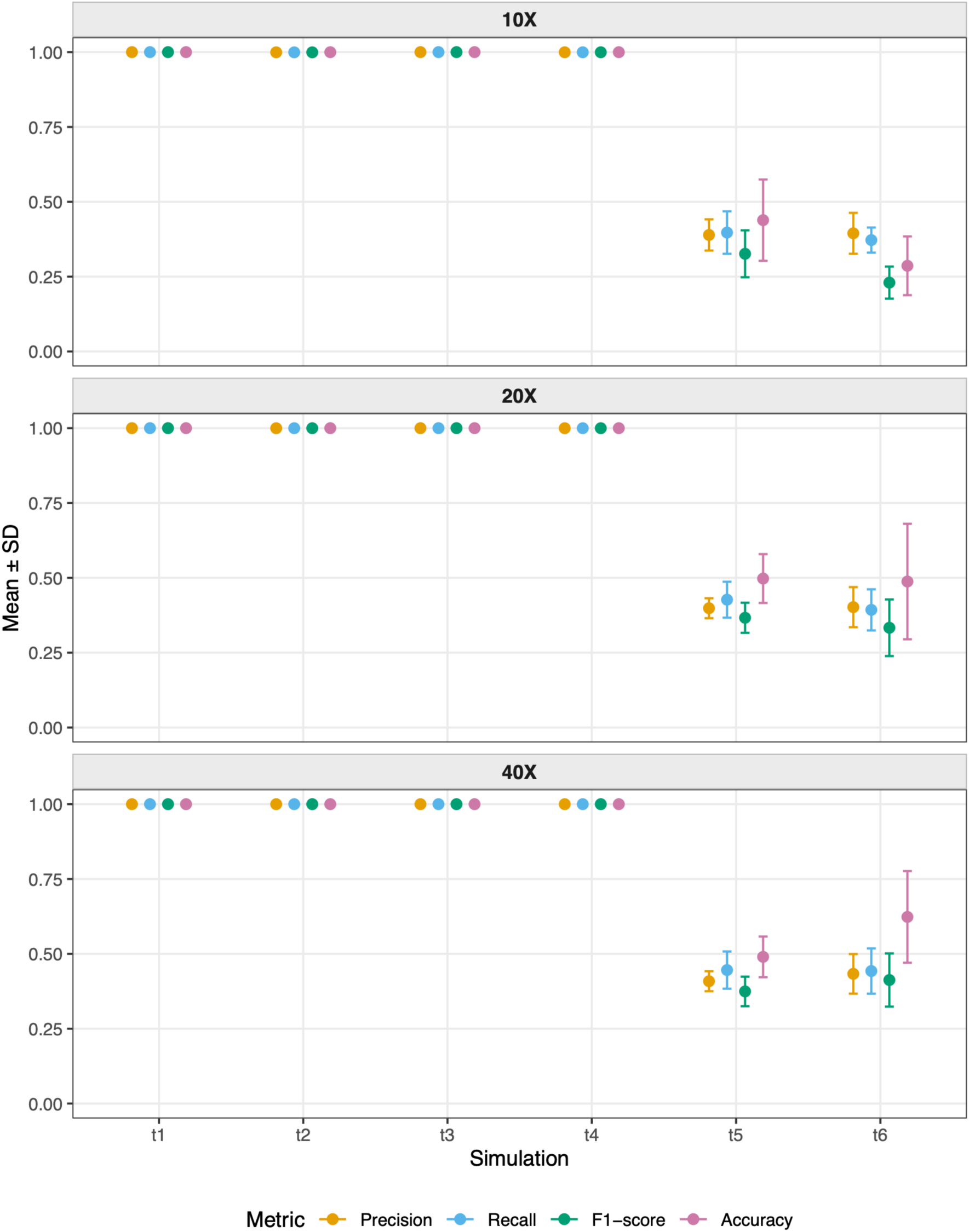
Logistic regression performance at variable coverages across simulations. Points are means and error bars are standard deviations. Simulations conditions on the x-axis refer to a *θ* of 0.0001 (t1), 0.0004 (t2), 0.001 (t3), 0.004 (t4), 0.01 (t5) and 0.04 (t6).

Results indicated that even low sequencing depths (e.g. 10x) should be sufficient for classification up to the ploidy levels explored here (Fig. 3; Fig. S7), as long as the level of divergence with a reference is appropriate. Because our simulations used only 1000 short loci, ploidy classification should be appropriate for Hyb-Seq or other sub-genomic approaches such as RADseq too. For similar-sized datasets, analysis with PPGTK *classify-ploidy* should be fast, taking at most 13 seconds on a single thread of an AMD Ryzen Threadripper for the largest simulated VCF. In the absence of a well-assembled reference genome, a *de novo* assembly of an outgroup or appropriate representative sample could be used. The effects of reference choice and performance on Hyb-Seq data are explored through analyses of empirical data below.

#### Empirical Analyses: Vaccinium sect. Cyanococcus

The true blueberries of North America (*Vaccinium* sect. *Cyanococcus* A. Gray) represent an ecologically and economically important polyploid complex of approximately 15 species. Despite their importance and long history of study, the taxonomy and evolutionary history of the group have remained enigmatic, partially due to poorly understood ploidy variation, which is not always morphologically detectable (but see Fritsch et al. 2026). Recent studies, however, have begun shedding light on blueberries and their wild relatives and have generated large phylogenomic datasets paired with flow cytometry-based ploidy estimates (e.g. Crowl et al. 2022; Fritsch et al. 2025; Manos et al. 2026). These data present a unique opportunity to validate our ploidy estimation method. We reanalyzed Hyb-Seq datasets (Angiosperms353 probes) from these studies, which include 70 individuals representing 15 named species, and one known artificially crossed F1 homoploid hybrid. FASTQ data were downloaded from NCBI SRA (Table S3), and a reference sequence was assembled for the outgroup, *V. macrocarpon*, using HybPiper (v2; Johnson et al. 2016) under default settings. We also explored the effects of using ingroup reference genomes. Specifically, we tested effects of the first haplotype of the haplotype-phased *V. darrowii* diploid reference genome (Yu et al. 2021) and the consensus of the highly mosaic tetraploid *V. corymbosum* genome (Colle et al. 2019). The analyses corresponding to the *V. macrocarpon*, *V. darrowii*, and *V. corymbosum* references are henceforth referred to as Vmac, Vdar, and Vcor, respectively.

Genotyping proceeded against the three references as follows: adapter removal with fastp v1.0.1 (Chen et al. 2018), alignment with BWA v0.7.17 using the MEM algorithm and returning only properly paired reads (-f 2), genotyping with HaplotypeCaller in GATK v4.6.2.0 (McKenna et al. 2010; DePristo et al. 2011; Poplin et al. 2017) in GVCF mode and GenotypeGVCFs from a GenomicsDB datastore, with a minimum genotype quality of 20 for calling. Hard-filtering of variants was then performed. First, the following GATK filter expressions were applied: FS > 60.0, MQ < 40.0, ReadPosRankSum < -8.0, MQRankSum < -12.5, and QD < 2.0. Filtered variants were excluded and only biallelic sites were retained. Then, vcftools v0.1.17 (Danecek et al. 2011) was used to filter variants with a depth less than 5 and sites with a mean depth less than 5. A second pass was then applied to reduce the VCF to biallelic sites only. The number of sites present after filtering was 51349, 1395670, and 696726 for Vmac, Vdar, and Vcor, respectively. Notably, there are large differences in the data based on mapping against targeted loci or whole genomes, and retaining some of this messiness was desired to evaluate robustness of the classification method. The PPGTK *classify-ploidy* logistic regression model was then applied to the Vmac, Vdar, and Vcor datasets under default settings, providing the known ploidy estimates of each individual to train and evaluate the model. Analyses with *classify-ploidy* took 16, 34, and 13 seconds for the Vcor, Vdar, and Vmac analyses, respectively, on a single thread of an Apple M4 processor.

A phylogeny was also generated to aid visualization and act as a sanity check on the genotyping results. A SNP-based maximum likelihood phylogeny from Hyb-Seq data is certainly not the best choice, given the known hybridization and high ILS in the group, but some taxonomic consistency should still be expected. The Vmac VCF was further filtered with vcftools to consecutively remove sites with 50%, 40%, 30%, and 20% missing data. No maximum depth filter was applied due to the high skew and inconsistency in coverage across target loci. This VCF of 49183 sites was converted to FASTA format with IUPAC codes for heterozygous sites. A total of 7576 biallelic variants (some sites considered biallelic in a VCF ultimately end up constant), of which 3387 were parsimony informative, were analyzed with IQTREE v3.1.2 (Wong et al., 2026) using ModelFinderPlus (Kalyaanamoorthy et al., 2017) across models with an ascertainment bias correction (Lewis, 2001) and 1000 ultrafast bootstraps (Hoang et al., 2018).

Overall, the ploidy classification performed very well where anticipated (Fig. 4). The Vmac and Vdar analyses predicted all samples correctly except one (SL410052_V_boreale_CY113). However, the taxonomic identification and original flow cytometry results for this individual are now called into question as a result of this study (Peter Fritsch pers. comm.). Otherwise, the Vcor analysis made three additional errors. Thus, there appears to be an effect of the underlying reference genome, with the outgroup and the targeted loci only yielding the sharpest kernel density estimates of the AB distribution (Fig. S8). These results are promising, as they imply analyses should be reasonably robust to reference choice that should be readily available for most Hyb-Seq-based biodiversity studies. Notably, the AB distributions across ploidies also demonstrate a reference bias in genotyping (Gerard 2018) that is especially prevalent in polyploids (Phillips 2024), such that the 0.25 and 0.75 AB ratios are never equal. These biases would likely lead to increasing uncertainty for probabilistic modeling (Tiley et al. 2024), but they are not problematic for the logistic regression approach. Some individuals do not appear where expected in the phylogeny, such as a rouge *V. corymbosum* or *V. tenellum* (Fig. 4), but overall nodes with decent support (ultrafast bootstrap > 0.95) recover clades of species or closely-related species. Therefore, the VCF needed for ploidy classification may have some analytical value beyond the classification itself.

**Figure 4.**
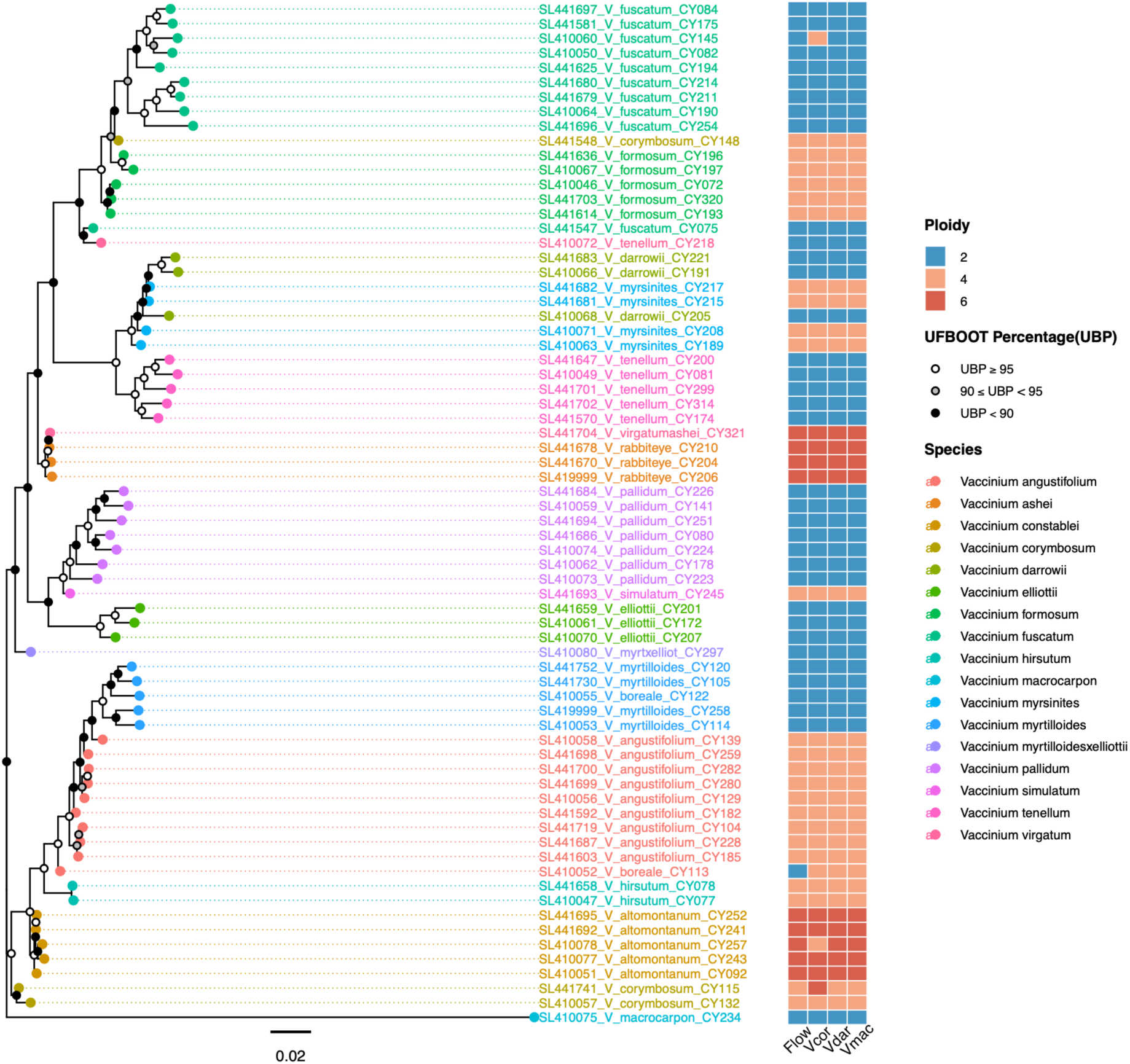
Maximum likelihood tree and ploidy classifications for *Vaccinium* sect. *Cyanoccocus*. Points at nodes are ultrafast bootstrap support values and the heatmap shows ploidy determined through flow cytometry or with classify-ploidy for the Vcor, Vdar, and Vmac analyses. This phylogenetic estimation was produced from a SNP matrix derived from Hyb-Seq data and is presented as a visualization aid for ploidy rather than a formal hypothesis of relationships. Note that tip labels are sample names from SRA based on initial determinations and do not reflect updated determinations as indicated in the legend.

#### Empirical Analyses: Ipomoea ser. Batatas

Although whole-genome methods exist for ploidy estimation (Ranallo-Benavidez et al. 2020; Rhie et al. 2020) there is no reason for the model to be limited to a specific data type, and we wanted to explore the potential for PPGTK to handle much larger VCFs. Additionally, whole-genomes could allow additional testing of filter effects. Thus, we reanalyzed 96 individuals representing seven species from *Ipomoea* ser. *Batatas* and a number of synthetic crosses (Table S4). The cultivated *I. batatas*, sweetpotato, is an important staple crop that, similar to blueberry, represents a plant breeding success despite persistent taxonomic and evolutionary uncertainty in the complex. Unlike the blueberry data though, the provenance of the known ploidy estimates is less known, and ploidy is accepted from the literature or encoded as “NA” if an individual could not be verified.

For the sweetpotato analyses, all individuals were genotyped using the diploid *I. triloba* genome as a reference (Wu et al. 2018). Otherwise, the genotyping steps were the same as our blueberry analyses. Different filtering choices were applied though. In vcftools, variants with a depth of less than 10 were dropped and sites with a mean depth of less than 10 were removed. A version of the VCF was retained at this point, referred to as the minimal filtering analysis, The consecutive missing site filters were then applied, but alternating with removing individuals with high missingness too. The procedure was: 1) remove sites over 50% missing data, 2) remove individuals over 90% missing data, 3) remove sites over 40% missing data, 4) remove individuals over 70% missing data, 5) remove sites over 30% missing data, and 6) remove individuals over 50% missing data. This procedure roughly follows O’Leary et al. (2018) and resulted in dropping 14 individuals from the initial sample [most were discarded by Yan et al. (2024) too]. The mean depth per site was then calculated and that grand mean plus two standard deviations in depth were used to create a max mean depth filter of 161x. Finally sites with more than 20% missing data were removed and this VCF was retained for analysis, called rigorous filtering. The minimal filtering analysis retained 356,651,033 sites and 96 individuals. The rigorous filtering retained 33,336,330 sites across 82 individuals. To make analyses comparable across filter sets only the 82 individuals in common were used for analysis with the logistic regression classifier. The *classify-ploidy* analysis took 21 minutes for the rigorously filtered VCF and 22 minutes for the minimally filtered VCF on a single thread of an Apple M4 processor. To generate a tree to aid with visualization, the rigorous filtering analysis was thinned to no more than one variant every 10,000 bp, which resulted in 40,111 sites. These sites were reduced to 8015 SNPs by IQTREE, such that 3326 SNPs were parsimony informative. Model selection and bootstrapping proceeded the same as the blueberry analyses in IQTREE.

Performance was overall high for the sweetpotato analyses, with an accuracy of 0.97 for the rigorous filtering that was reduced only slightly to 0.94 with minimal filtering. One classification error shared by both filtering strategies called an *I. aequatoriensis* individual (SRR17610138) as diploid (Fig 5). All *I. aequatoriensis* are presumably tetraploid, but this individual does not fall in the otherwise well-supported (ultrafast bootstrap > 0.95) *I. aequatoriensis* and instead forms a clade with other diploid more-distant relatives of sweetpotato. Thus, this classification error may represent a misidentification or errors in the underlying metadata. The other error for the rigorous filtering involved calling a tetraploid *I. batatas* individual (CRR284435) as hexaploid.

**Figure 5.**
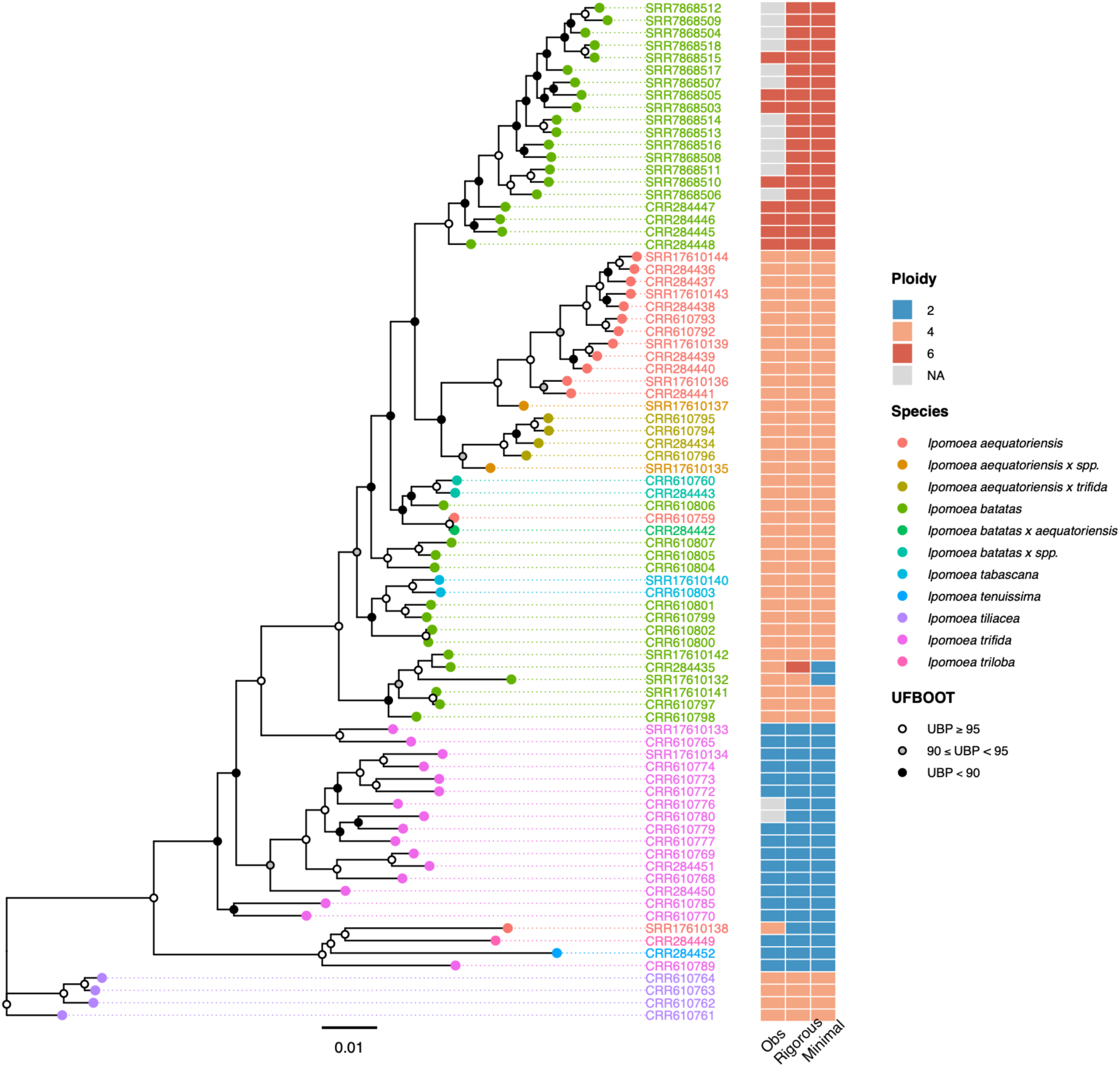
Maximum likelihood tree and ploidy classifications for *Ipomoea* ser. *Batatas*. Points at nodes are ultrafast bootstrap support values and the heatmap shows ploidy determined through prior analyses or with *classify-ploidy* for the rigorous and minimal filtering strategies. This phylogenetic estimation is provided- as a visualization aid for ploidy rather than a formal hypothesis of relationships. Individuals without prior information are coded as “NA” in the input metadata and not utilized in model training.

This likely reflects a legitimate mis-classification error as hexaploid versus tetraploid *I. batatas* tend to form clades (Fig. 5), but the mis-classification is understandable as the AB distribution looks much more similar to other hexaploids rather than tetraploids (Fig. S9). In a more-focused organismal study, this individual would warrant inspection to understand other properties of the sequence data or genotyping process driving the AB distribution. Otherwise, the F1-Score was 0.97, 0.98, and 0.94 for diploids, tetraploids, and hexaploids, respectively with rigorous filtering. This reflects typically acceptable recall and precision, and there was only a slight decrease in light of likely egregious false positives in the VCF, for which the minimal filtering is a proxy (Table 1). For the unknown *I. batatas* and *I. trifida* individuals in the sample, all were classified as hexaploid or diploid as expected based on their position in the tree with both the rigorous and minimal filtering VCFs (Fig. 5).

**Table 1.**
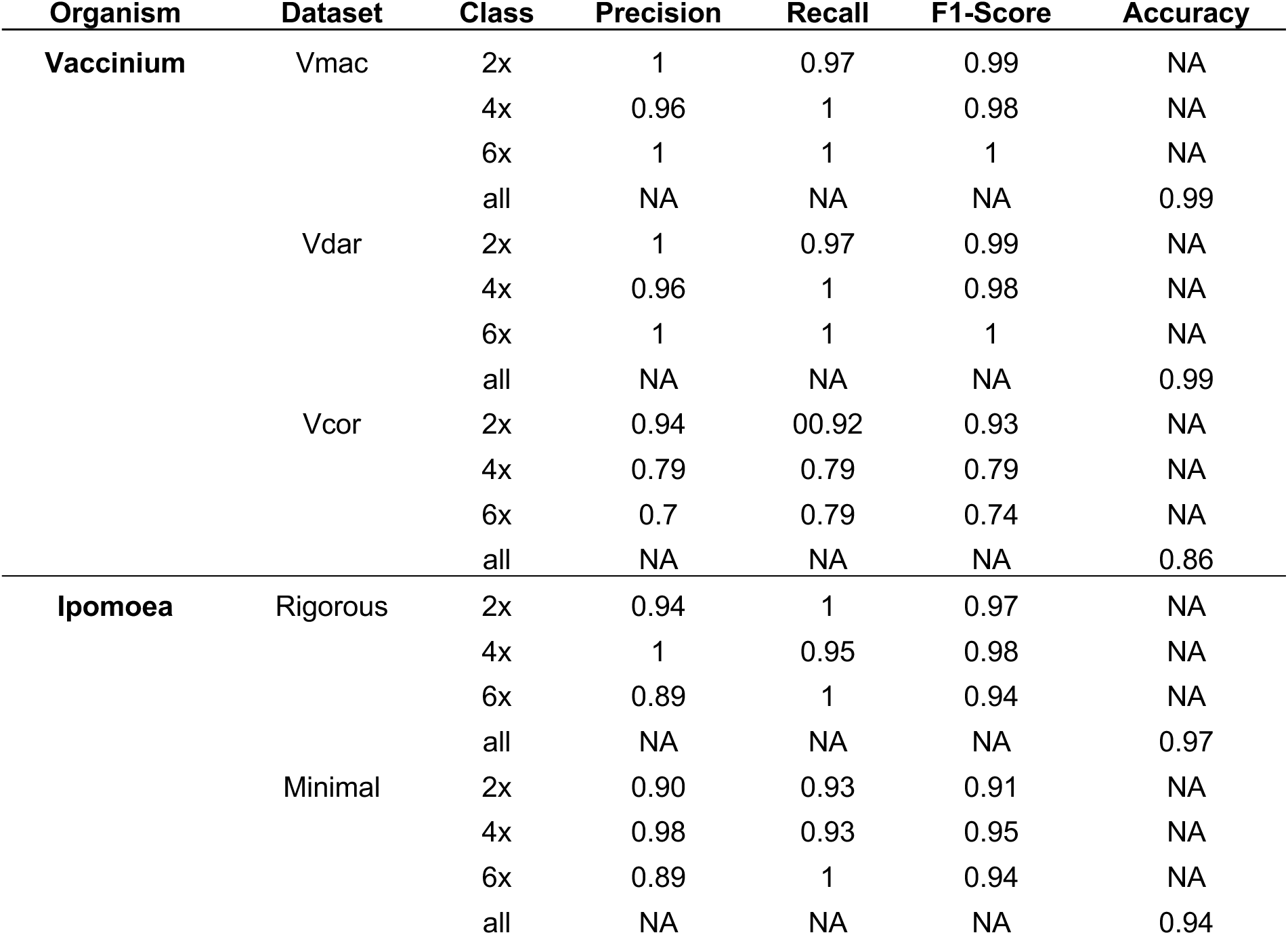
Performance statistics on empirical data.

## CONCLUSIONS

We found the logistic regression ploidy classification in PPGTK (*classify-ploidy)* to be a simple and promising approach to recover ploidy directly from sequence data. Simulations suggested 0.99 or greater accuracy could be expected for systems where reads are still alignable to a reference with as low as 10x coverage. Our analyses of empirical datasets resulted in overall accuracies of 0.99 across 70 *Vaccinium* individuals and 0.97 across 82 *Ipomoea* individuals, and that reliable results could be obtained from diverse datatypes (Table 1). The method requires only sequence data in VCF format and a table of known ploidy levels, making it ideal for projects in which some samples have been processed with flow cytometry or chromosome counts.

A few caveats should be considered. The ploidy prediction itself is efficient, though we are hesitant to call the entire process fast, as the initial genotyping procedure can be computationally and time intensive depending on the number of individuals and amount of data. If a multisample VCF is needed for routine population genomic analyses though, our approach becomes advantageous. Genotyping polyploid individuals as diploids can still be useful as some analyses are insensitive to allele dosage (e.g. Meirmans et al. 2024), but a study could build upon a diploid-genotyped VCF and our classifications to enable polyploid genotyping across all samples (Blischak et al. 2018; Gerard et al. 2018; Clark et al. 2019). The taxonomic breadth considered by our approach has limitations, since reference quality and polarization of alleles had a larger effect than sequencing depth (Figs. 3 and 4), but in such a scenario, breaking a larger tree into individual clades should be feasible for a Hyb-Seq study.

A potential complication not addressed here is when some ploidies are unobserved in the training data. Training data ideally has representation from every ploidy level possible in the unknown samples, as the classifier cannot predict a class it has never seen. For example, if data are only trained on diploids, tetraploids, and hexaploids, it will not be feasible to correctly identify a triploid or pentaploid. In such cases, the per-class probabilities should be considered. We anticipate individuals not fitting an *a priori* ploidy class would propagate uncertainty with non-negligible weight among multiple classes. An assessment of these individual probability outputs from PPGTK *classify-ploidy* would allow a researcher to make informed judgement calls on the reliability of results.

We expect our findings and the PPGTK application should be helpful for the analysis of museum specimens, where flow cytometry is not possible, or large-scale population studies where ploidy estimation in the lab for thousands of individuals is not feasible. PPGTK *classify- ploidy* should make it possible to leverage ploidy information from a subset of individuals to address questions about polyploid evolution both within and between species.

## Author Contributions

GPT and AAC conceived the study. SVK and GPT developed code and analyzed data. GPT developed the simulation pipeline. GPT, SVK, and AAC wrote the first draft. All authors revised and approved the submitted manuscript.

## Acknowledgements

Startup funds from NCSU contributed to this research. The authors thank Peter Fritsch and Paul Manos for helpful discussions on *Vaccinium*.

## Data Availability Statement

VCFs and metadata needed to reproduce *Vaccinium* sect. *Cyanococcus* and *Ipomoea* ser. *Batatas* analyses with PPGTK will be made available via Dryad upon acceptance. PPGTK is available on GitHub, and release v0.1.0-alpha was the version used for analyses in this manuscript (https://github.com/tileylab/PPGTK/releases/tag/v0.1.0-alpha). PPGTK currently has other functions for calculating population genetic summary statistics, but the *classify-ploidy* function implements the machine-learning method described in the manuscript.

