## Supplementary Material for "A simple and accurate method for inferring missing ploidy information from sequence data"

### Supplementary Figures

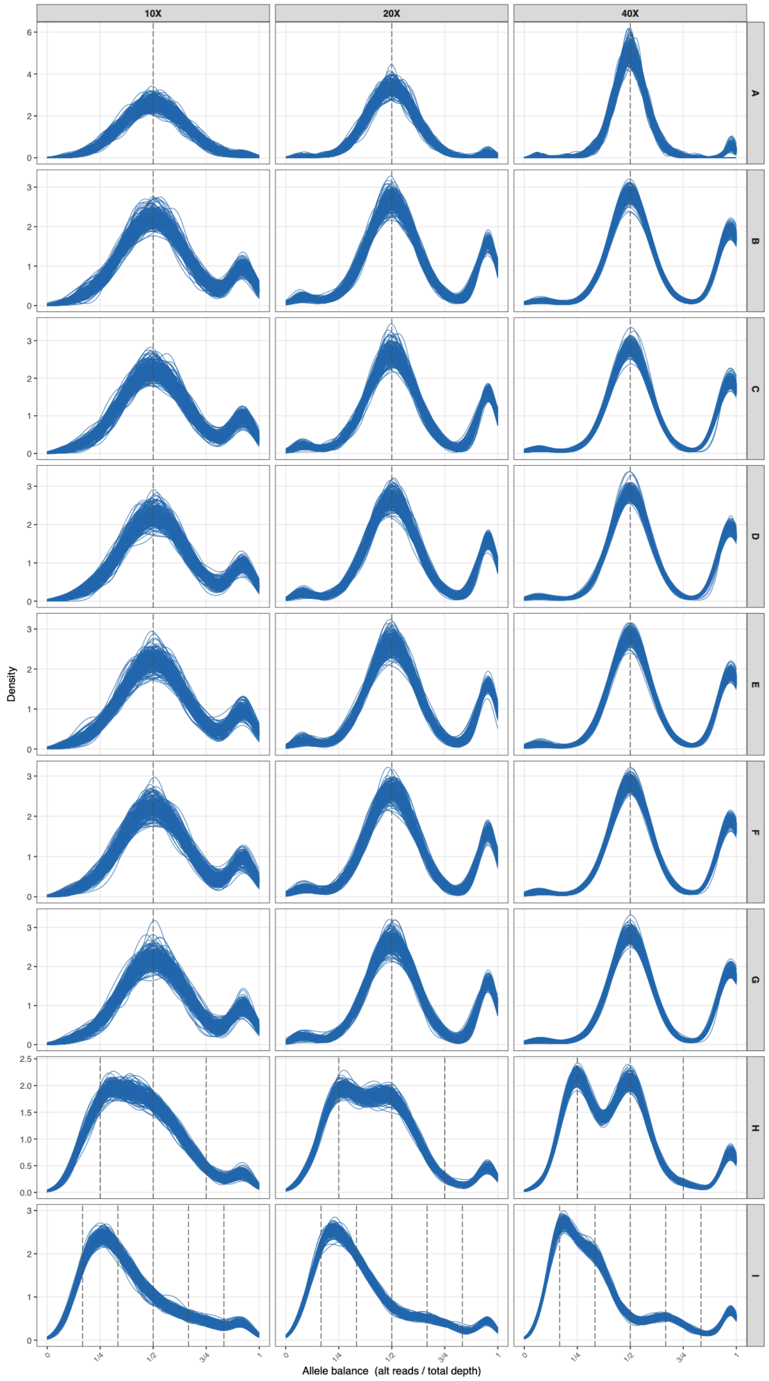


**Figure S1 — Allele balance distribution for simulations with $\tau_t = 0.001$ and $\theta = 0.001$.** Distributions are shown for 20 individuals across 10 simulation replicates.

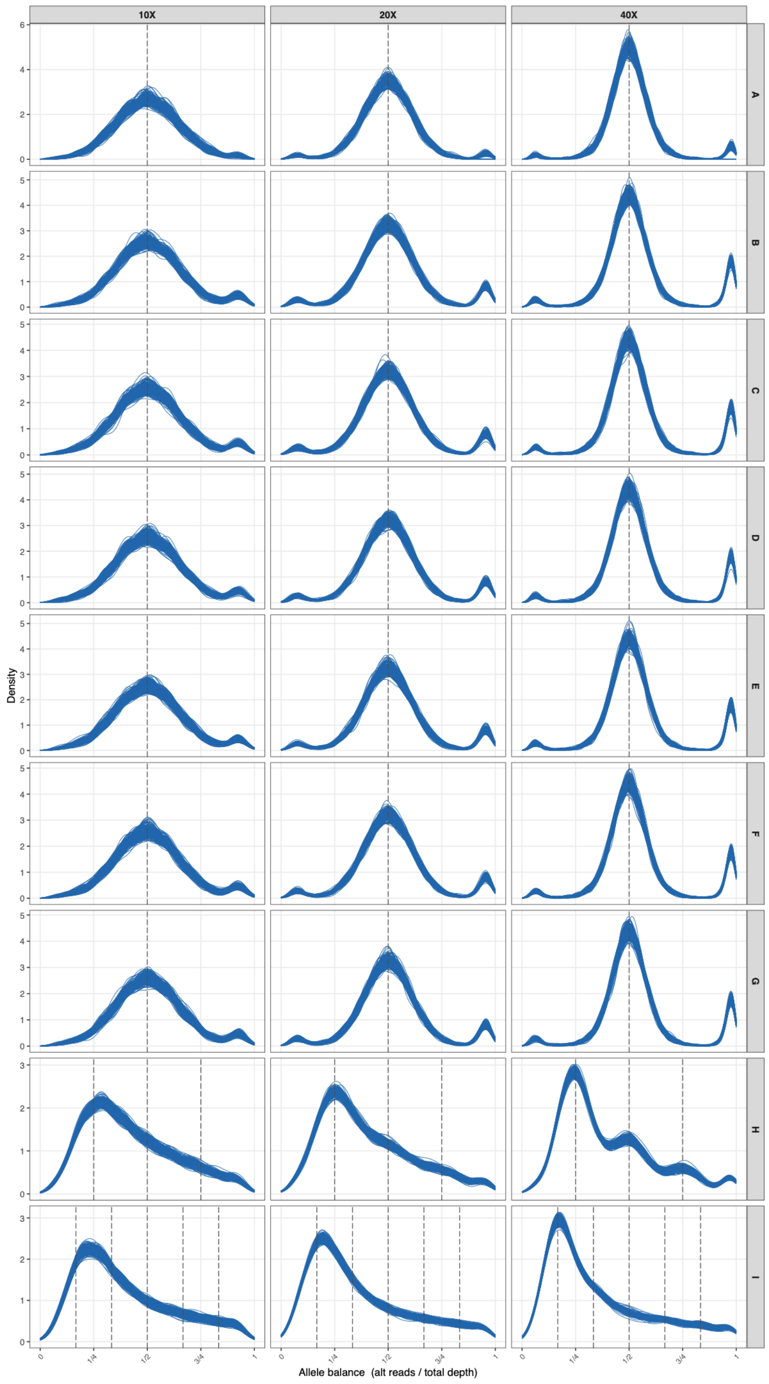


**Figure S2 — Allele balance distribution for simulations with $\tau_t = 0.001$ and $\theta = 0.004$.** Distributions are shown for 20 individuals across 10 simulation replicates.

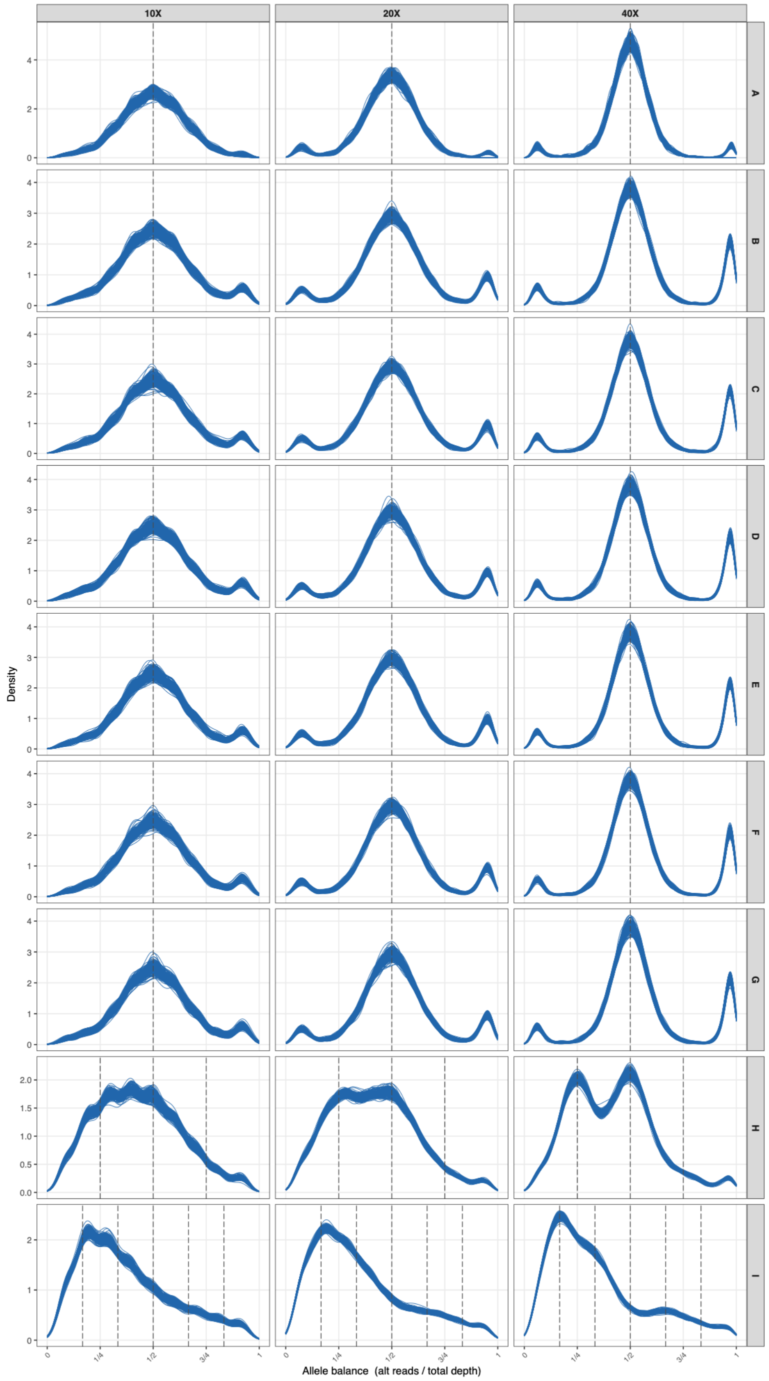


**Figure S3 — Allele balance distribution for simulations with $\tau_t = 0.01$ and $\theta = 0.01$.** Distributions are shown for 20 individuals across 10 simulation replicates.

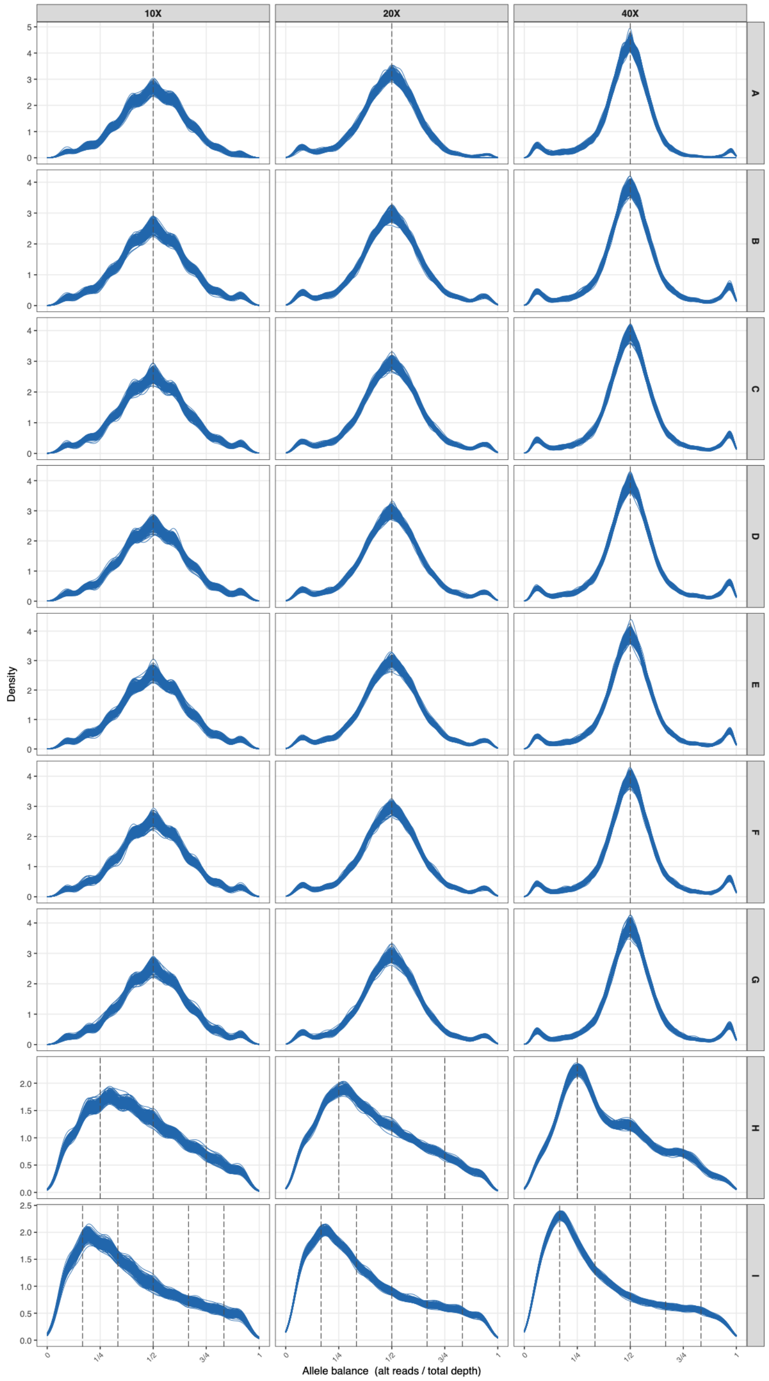


**Figure S4 — Allele balance distribution for simulations with $\tau_t = 0.01$ and $\theta = 0.04$.**
Distributions are shown for 20 individuals across 10 simulation replicates.

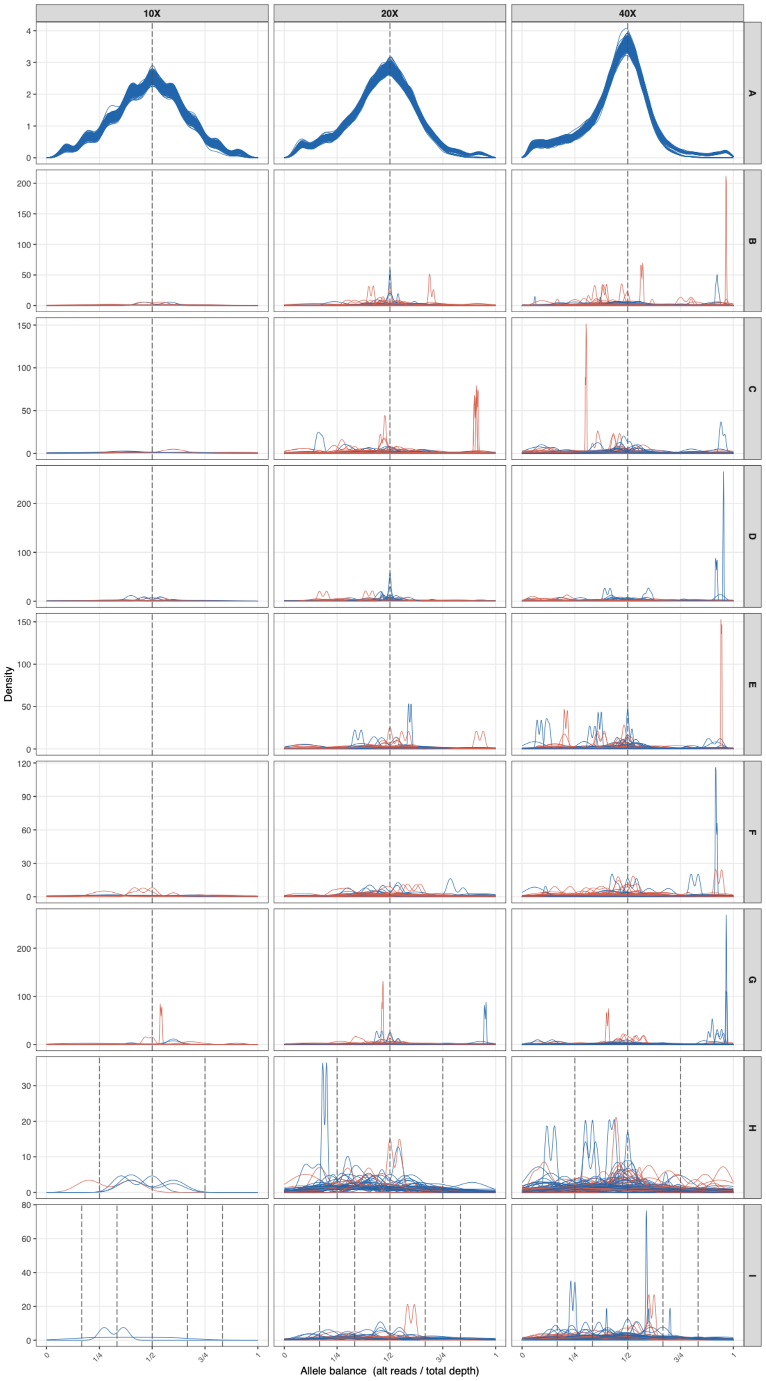


**Figure S5 — Allele balance distribution for simulations with $\tau_t = 0.1$ and $\theta = 0.1$.** Distributions are shown for 20 individuals across 10 simulation replicates. Blue lines indicate correctly classified individuals and red lines indicate incorrectly classified individuals in subsequent logistic regression analyses.

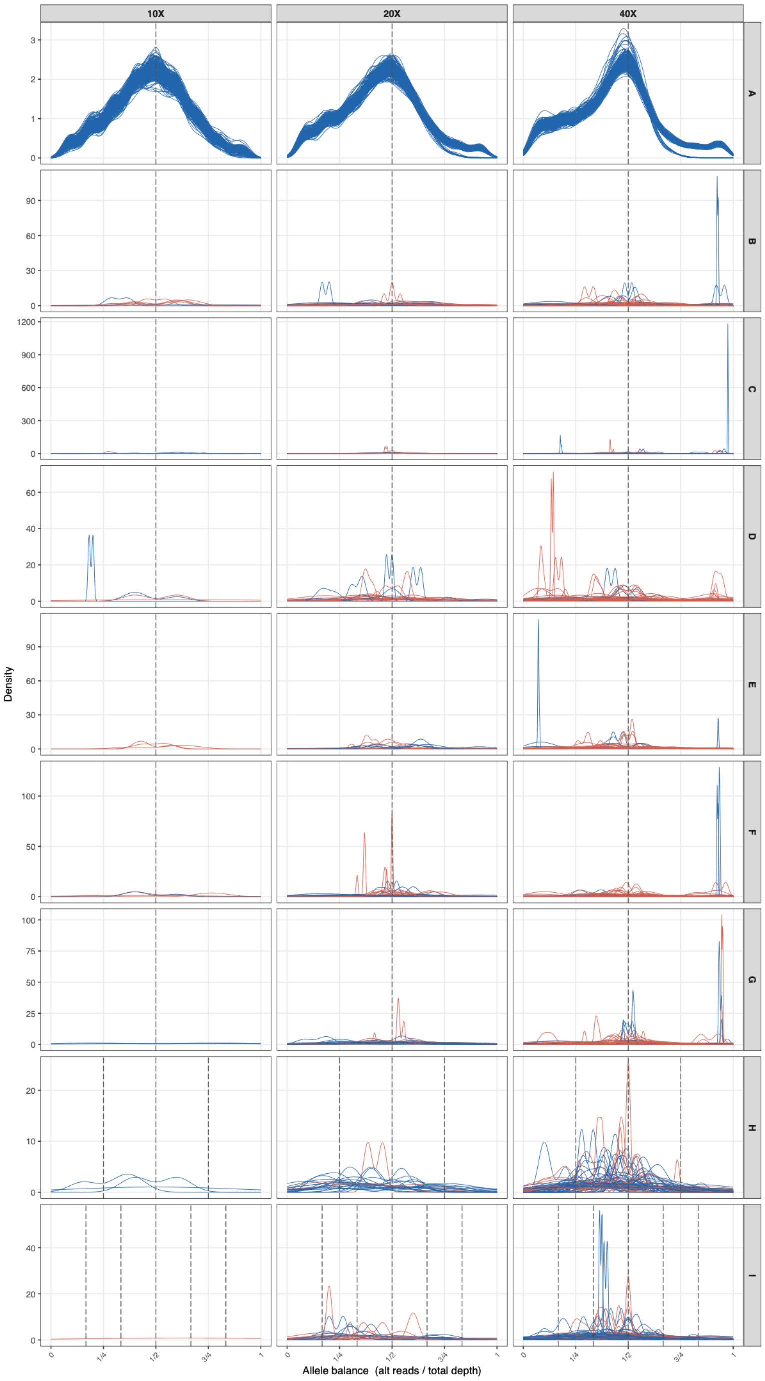


**Figure S6 — Allele balance distribution for simulations with $\tau_t = 0.1$ and $\theta = 0.4$.** Distributions are shown for 20 individuals across 10 simulation replicates. Blue lines indicate correctly classified individuals and red lines indicate incorrectly classified individuals in subsequent logistic regression analyses.

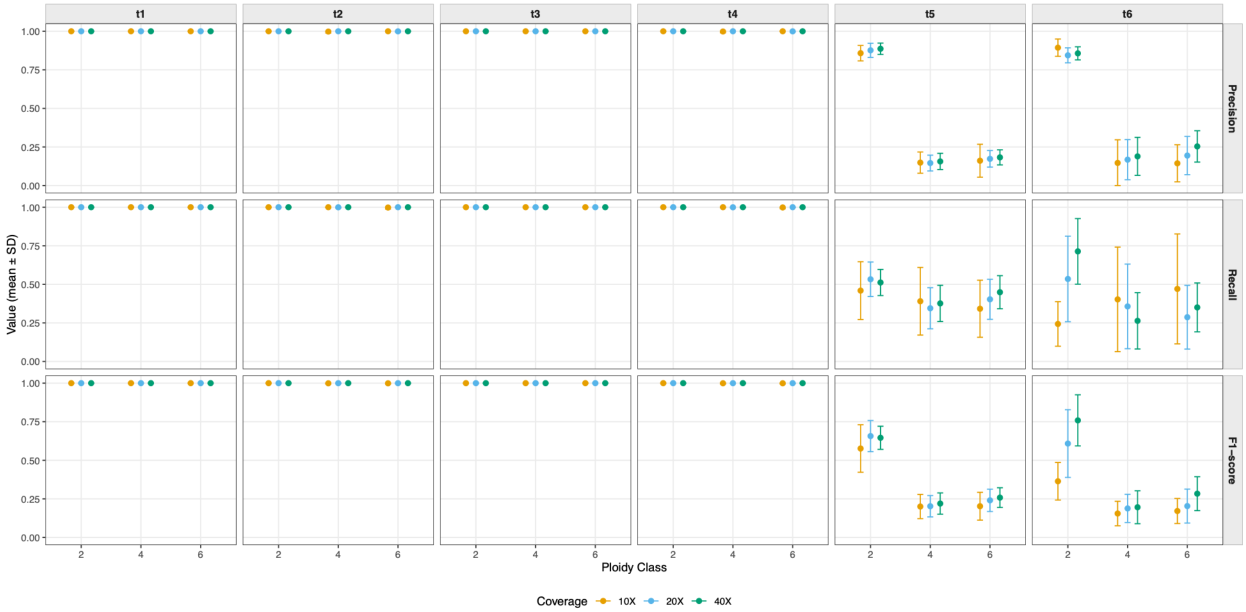


**Figure S7 — Per-class metrics across simulation scenarios and depths.** Means and standard deviations are calculated from 100 replicates.

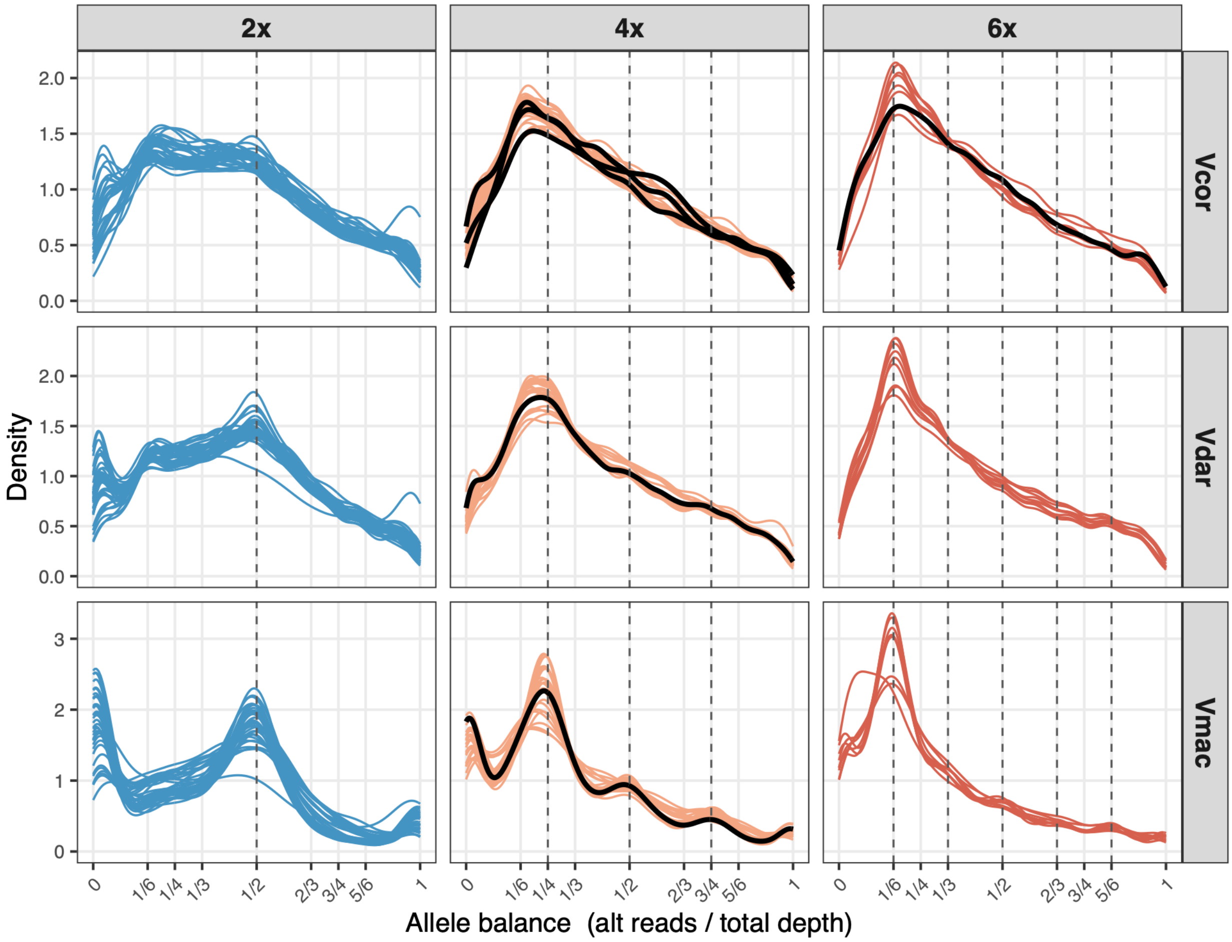


**Figure S8 — Allele balance distributions for *Vaccinium* spp. individuals by ploidy across analyses.** Black lines show incorrectly classified individuals. One individual (SL410052_V_boreale_CY113) was scored as incorrect across analyses as the flow cytometry data originally indicated it was diploid, but the distinct peak around 0.25 is characteristic of other tetraploids.

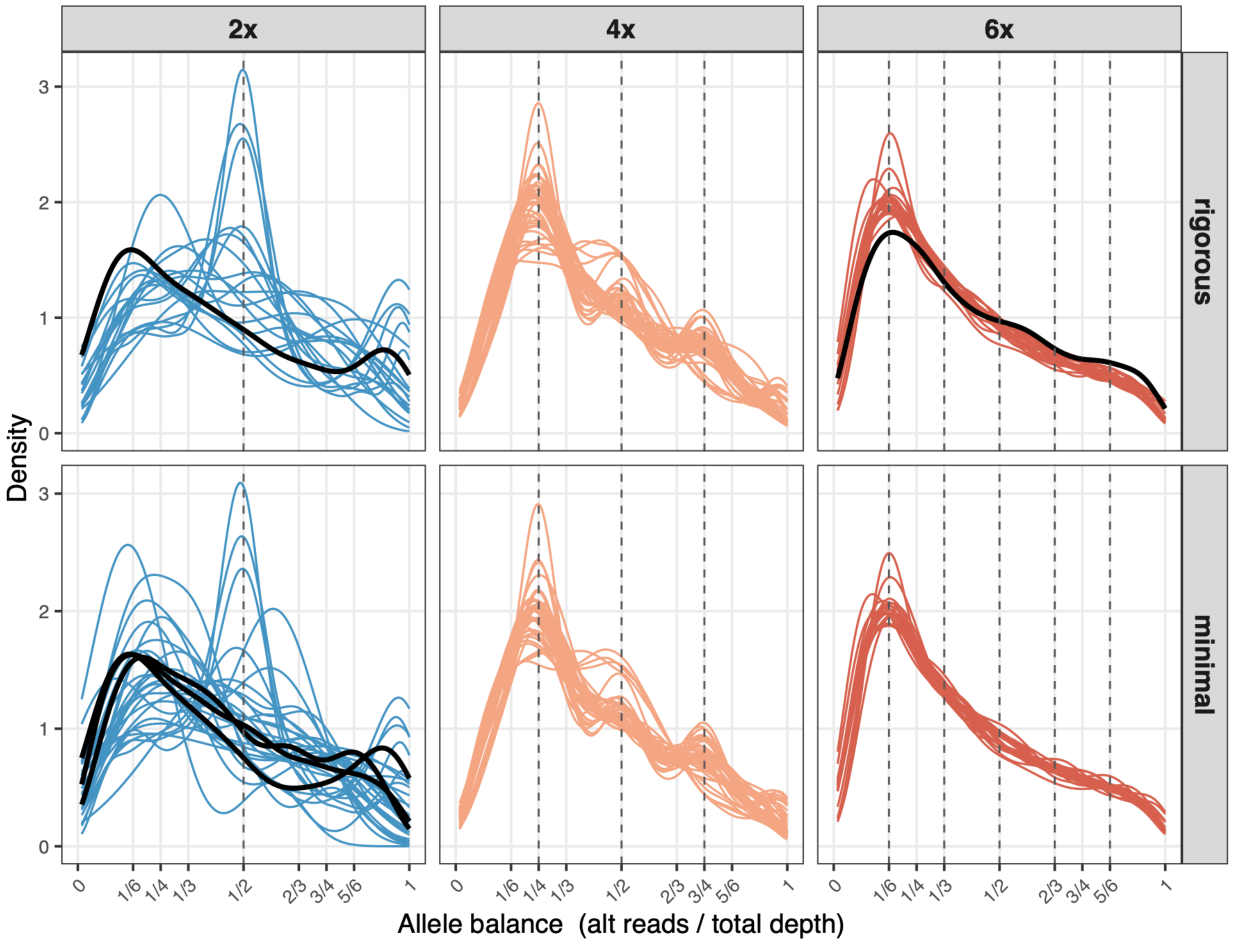


**Figure S9 — Allele balance distributions for *Ipomoea* spp. individuals by ploidy across analyses.** Black lines show incorrectly classified individuals. One diploid individual (SRR17610138) was scored as incorrect across both analyses, but the placement of an *I. aequatoriensis* individual in a well-supported clade with earlier-diverging individuals is suspect. Another tetraploid individual was scored incorrectly in both analyses, but the rigorous filtering classified it as a hexaploid while minimal filtering classified it as a diploid.
